# Plxdc family members are novel receptors for the rhesus monkey rhadinovirus (RRV)

**DOI:** 10.1101/2020.01.20.912246

**Authors:** Anna K. Großkopf, Sarah Schlagowski, Thomas Fricke, Armin Ensser, Ronald C. Desrosiers, Alexander S. Hahn

**Affiliations:** German Primate Center - Leibniz Institute for Primate Research, Göttingen, Germany; Universitätsklinikum Erlangen, Institute for Clinical and Molecular Virology, Erlangen, Germany; Miller School of Medicine, University of Miami, Miami, United States

## Abstract

The rhesus monkey rhadinovirus (RRV), a γ2-herpesvirus of rhesus macaques, shares many biological features with the human pathogenic Kaposi’s sarcoma-associated herpesvirus (KSHV). Both viruses, as well as the more distantly related Epstein-Barr virus, engage cellular receptors from the Eph family of receptor tyrosine kinases (Ephs). However, the importance of the Eph interaction for RRV entry varies between cell types suggesting the existence of Eph-independent entry pathways. We therefore aimed to identify additional cellular receptors for RRV by affinity enrichment and mass spectrometry. We identified an additional receptor family, the Plexin domain containing proteins 1 and 2 (Plxdc1/2) that bind the RRV gH/gL glycoprotein complex. Preincubation of RRV with soluble Plxdc2 decoy receptor reduced infection by approx. 60%, while overexpression of Plxdc1 and 2 dramatically enhanced RRV susceptibility of otherwise marginally permissive Raji cells. While the Plxdc2 interaction is conserved between two RRV strains, 26-95 and 17577, Plxdc1 specifically interacts with RRV 26-95 gH. The Plxdc interaction is mediated by a short motif at the N-terminus of RRV gH that is partially conserved between isolate 26-95 and isolate 17577, but absent in KSHV gH. Mutation of this motif abrogated the interaction with Plxdc1/2 in *in vitro* assays and reduced RRV infection in a cell-type specific manner. Taken together, our findings characterize Plxdc1/2 as novel interaction partners and entry receptors for RRV and support the concept of the N-terminal domain of the gammaherpesviral gH/gL complex as a multifunctional receptor-binding domain. Further, Plxdc1/2 usage defines an important biological difference between KSHV and RRV.

**AUTHORS SUMMARY:** KSHV is the causative agent of a group of malignancies which account for a substantial disease burden in particular in sub-Saharan Africa. RRV, a related virus of rhesus macaques, has shown promise as model system for KSHV and for the development of immunization strategies. To exploit the full potential of the RRV animal model system, detailed knowledge of commonalities and differences between KSHV and RRV is key. Here, we describe the Plexin domain containing proteins 1 and 2 as a novel receptor family which mediates entry of RRV, but not of KSHV. Infection experiments using RRV mutants deleted of the Plxdc interaction motif suggest a cell type-specific contribution of Plxdc receptors to RRV infection. As information on Plxdc1/2 and its biological function is still sparse, analysis of the RRV–Plxdc interaction will help to characterize the physiological and pathophysiological role of this receptor family.

## INTRODUCTION

The rhesus monkey rhadinovirus (RRV), a member of the genus γ2-herpesvirus or rhadinovirus, is closely related to the only human pathogenic member of this genus, the Kaposi’s sarcoma-associated herpesvirus (KSHV) (1, 2). Due to the high similarity in both genome organization and biology RRV is considered as an animal model virus for KSHV (3) and has been used as such in *in vitro* and *in vivo* studies. Two major RRV sequence groups have been identified (4), and each is represented by a cloned isolate, RRV 26-95 (5) and RRV 17577 (6). Analogous to KSHV infection, primary RRV infection is asymptomatic in healthy hosts and leads to life-long persistence, most likely in the B cell compartment (7). KSHV is associated with a solid tumor of endothelial origin, Kaposi’s sarcoma (KS), and two B cell malignancies, primary effusion lymphoma (PEL) and the plasmablastic variant of multicentric Castleman’s disease (MCD), most prominently in the context of human immunodeficiency virus (HIV) infection and in immunocompromised individuals. Similarly, simian immunodeficiency virus (SIV)-positive rhesus macaques developed B cell lymphomas upon experimental infection with RRV strain 17577 (8, 9) and several studies correlated RRV infection with lymphomagenesis in SIV/SHIV-infected animals (10, 11). While RRV is not consistently associated with solid malignancies, RRV has been identified in retroperitoneal fibromatosis tissue (9, 12), similar to retroperitoneal fibromatosis herpesvirus (RFHV)(11), and was recently isolated from hemangioma tissue (13). Another shared characteristic of KSHV and RRV is the receptor usage on a range of cell types. Both viruses engage members of the Eph family of receptor tyrosine kinases (Ephs) through their glycoprotein (g)H/gL complex to facilitate entry into target cells. While KSHV preferentially interacts with A-type Ephs – specifically EphA2 as the high affinity receptor (14, 15) – RRV can utilize both A- and B-type Ephs (15) for entry. These interactions have been characterized on different adherent cells types (6–9) and we could recently show that both viruses can utilize EphA7 as receptor on BJAB cells (20), a model B lymphocyte line. While for KSHV, in addition to Eph family receptors, several membrane proteins have been proposed as cellular receptors for different viral glycoproteins mediating either attachment or entry on a range of target cells (reviewed in (21)) the receptor usage of RRV is comparatively less well characterized. Nevertheless, studies using receptor knock-down or knockout (14, 22), receptor- and ligand-mediated blocking (15, 23), and Eph de-targeted virus mutants (23, 24) showed that both viruses and in particular RRV can infect various cells partially or completely independently of the Eph-interaction, which suggests that RRV engages at least one additional entry receptor that can functionally substitute for the Eph-interaction. This notion is also supported by a recent *in vivo* study that demonstrated that an RRV mutant deleted of gL and therefore unable to interact with Eph receptors still establishes persistent infection in RRV-naïve rhesus macaques upon intravenous inoculation (24). We therefore aimed to identify additional rhadinovirus receptors that bind the gH/gL complex or gH and identified Plexin domain containing protein 2 (Plxdc2) as novel interaction partner of the gH/gL complex of RRV, but not KSHV. The closest homolog to Plxdc2, Plxdc1 was initially identified as overexpressed in blood vessels of solid human tumors (25), resulting in the original terminology tumor endothelial marker 7 (TEM7, Plxdc1) and tumor endothelial marker 7 related (TEM7R, Plxdc2) (26). In general, the physiological functions of Plxdc1/2 are not well understood. Suggestive of a role in development, Plxdc2 has been described as mitogen for neural progenitor cells (27) and expression of both Plxdc1 and Plxdc2 in the developing nervous system has been demonstrated (28, 29). Cortactin, nidogen and the pigment epithelium derived factor (PEDF) have been described as interactors for Plxdc1 and Plxdc2 (30–32). However, the physiological relevance of these interactions is not fully understood. In this study we characterize the interaction of Plxdc1/2 with the gH/gL glycoprotein complex of RRV and establish Plxdcs as novel cellular RRV entry receptors.

## RESULTS

To identify potential cellular receptors for RRV glycoprotein H, we performed immunoprecipitation using soluble RRV 26-95 gH, consisting of the extracellular part fused to the Fc part of human IgG (RRV gH-FcStrep) as bait and 293T whole cell lysate as prey (Fig 1A). Bands present in the precipitation from 293T whole cell lysate, but not in control precipitation without 293T lysate were excised and analyzed by LC-MS/MS. The most abundant cell surface protein, identified in four of the five analyzed regions, was Plxdc2 or TEM7R, a cellular transmembrane protein. As described above, Plxdc1 or TEM7 is the only homolog of Plxdc2 in humans and rhesus macaques, and was therefore included in subsequent analyses. Human and rhesus Plxdc1 (ref |NM_020405.5|; ref |XM_028836436.1|) and Plxdc2 (ref |NM_032812.9|; ref |XM_028826043.1|) are 96.80% and 97.92% identical on the amino sequence level.

**Figure 1.**
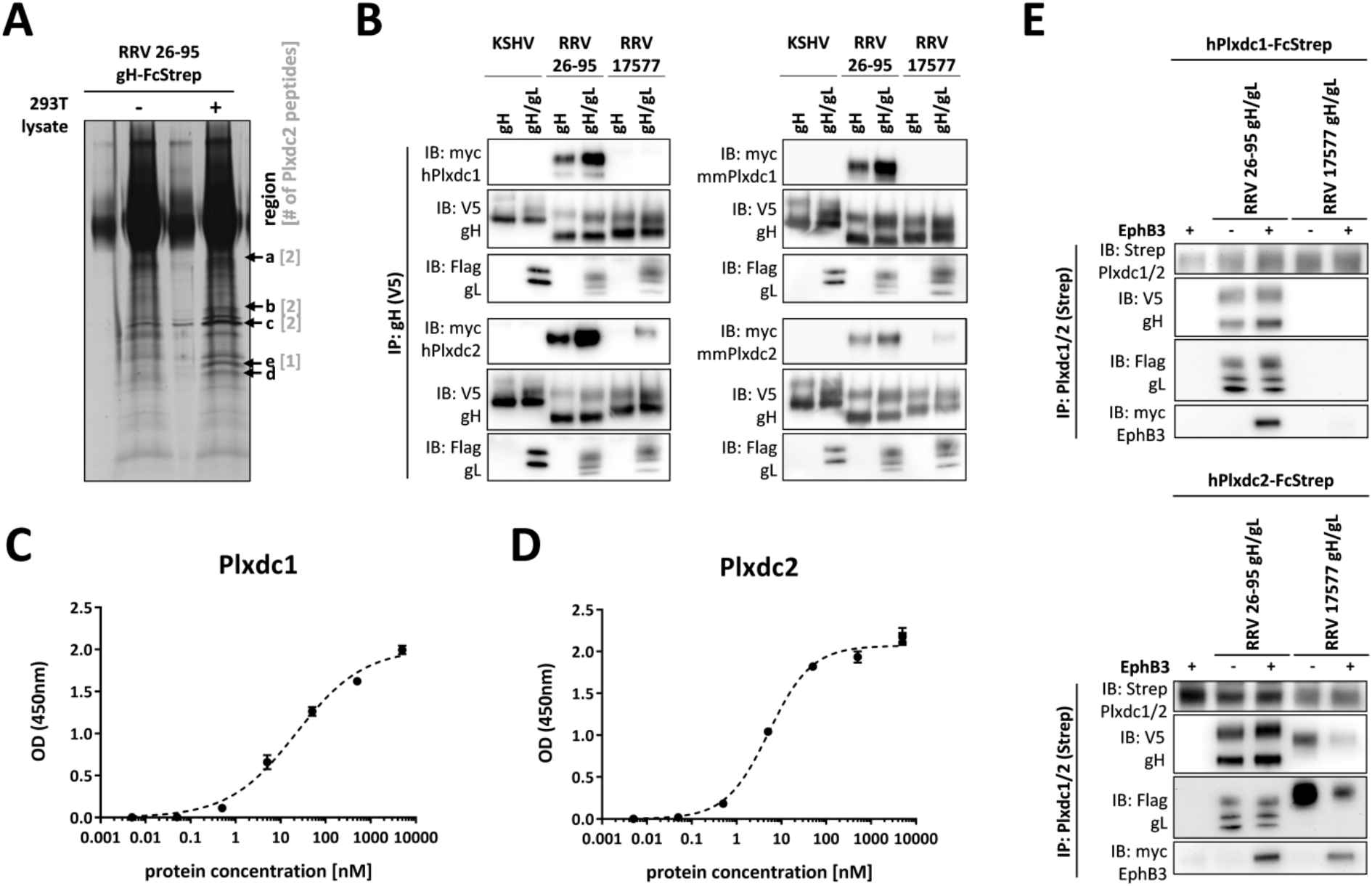
Plxdc family receptors are novel interaction partners for the RRV gH/gL. **A)** Immunoprecipitation of recombinant soluble RRV 26-95 gH-FcStrep in the presence or absence of 293T lysate. Precipitates were analyzed by PAGE, silver stained and bands at the indicated molecular weight (arrows, regions a-d) were excised and analyzed by mass spectrometry. Numbers in brackets indicate the number of Plxdc2 peptides identified by LC-MS/MS in each region. **B)** V5-tagged RRV 26-95 gH, RRV 17577 gH or KSHV gH alone or co-expressed with the respective Flag-tagged gL construct were immunoprecipitated in the presence of full-length Plxdc1 or Plxdc2 from human (h) or macaca mulatta (mm) origin using monoclonal antibody to the V5-tag. Precipitates were analyzed by Western blot with the indicated antibodies. **C-D)** Binding of Plxdc1 ectodomain C) or Plxdc2 ectodomain D) at various concentrations to immobilized RRV 26-95 gH-FcStrep/gL was measured by enzyme-linked immunosorbent assay. Curve Fitting and determination of half-maximal binding concentration was performed based on the *One site specific binding with Hill Slope equation* in Prism6. **E)** Co-immunoprecipitation of soluble human Plxdc1-FcStrep or human Plxdc2-FcStrep with RRV 26-95 gH-V5/gL-Flag or RRV 17577 gH-V5/gL-Flag using StrepTactin Sepharose in the presence or absence of human full-length EphB3. Abbreviations: IP: immunoprecipitation, IB: immunoblotting, h: human, mm: macaca mulatta (rhesus macaque).

Co-immunoprecipitation of V5-tagged expression constructs of gH from KSHV and from the two RRV isolates 26-95 and 17577, in the presence or absence of the corresponding Flag-tagged gL proteins with myc-tagged human Plxdc1 or Plxdc2 (hPlxdc1-myc/ hPlxdc2-myc) from transfected 293T cells confirmed the interaction of both RRV gH/gL complexes with Plxdc2 (Fig 1B). Neither KSHV gH/gL nor RRV 17577 gH/gL interacted detectably with Plxdc1-myc, while RRV 26-95 prominently bound both Plxdc1 and Plxdc2 in the presence and absence of gL. Measurement of the binding of purified Plxdc1 and Plxdc2 ectodomain to immobilized RRV 26-95 gH-FcStrep/gL by enzyme-linked immunosorbent assay (Fig 1C, D) showed a half-maximal binding concentration of 22.2nM for Plxdc1 and 5.2nM for Plxdc2 as calculated for one site specific binding without pre-defined Hill slope.

To evaluate the effect of Plxdc-binding to RRV gH on the interaction with EphB3, the high-affinity Eph family receptor for RRV gH/gL (15), we used soluble human Plxdc1 or Plxdc2, consisting of the extracellular part of Plxdc1 or Plxdc2 fused to the Fc part of human IgG followed by a TwinStrep tag (hPlxdc1-FcStrep/ hPlxdc2-FcStrep) in immunoprecipitation experiments. Co-immunoprecipitation of hPlxdc1-FcStrep/ hPlxdc2-FcStrep with the gH-V5/gL-Flag complexes of RRV isolates 26-95 and 17577 in the presence or absence of myc-tagged EphB3 from transfected 293T cells demonstrated the existence of a quaternary complex, indicating the ability of RRV gH/gL to interact with members of both receptor families simultaneously (Fig 1E).

While interaction of purified proteins and in transfected cell lysates is strongly suggestive of a functional interaction, the biologically relevant interaction for the entry process would occur with virion gH/gL. To evaluate the functionality of the gH/gL-Plxdc interaction on virus particles both for wildtype RRV and an Eph-binding-negative RRV mutant, we utilized RRV-YFP, an RRV 26-95 strain engineered for constitutive YFP expression upon infection, and RRV-YFP gH-AELAAN, an Eph-binding-negative RRV-YFP mutant that we had previously described (Fig 2A) (23). To analyze the impact of competition with soluble Plxdc decoy receptor, RRV-YFP wt and RRV-YFP gH-AELAAN preparations were incubated with a concentration series of soluble hPlxdc2-FcStrep or an FcStrep control prior to infection of HaCaT cells (Fig 2B). According to RNA-Seq data of 36 cell lines (Courtesy of Human Protein Atlas, www.proteinatlas.org, (33)), HaCaT cells exhibit the highest cell-line specific expression of Plxdc2 among the analyzed non-cancer cell lines and were therefore chosen for further analyses. Soluble Plxdc2-FcStrep inhibited RRV-YFP wt infection up to approx. 60% in a dose dependent manner when compared to FcStrep alone. Likewise, preincubation of RRV-YFP gH-AELAAN with hPlxdc2-FcStrep reduced infection by approx. 65%. RRV-YFP wt and RRV-YFP gH-AELAAN infection was normalized to approx. MOI 0.2. Inhibition of the gH/gL-Eph interaction, which served as control, lead to an approx. 50% reduction of RRV-YFP wt infection at a concentration of 10nM hEphB3-Fc while 100nM of soluble hPlxdc2-FcStrep exhibited a similar blocking efficiency. Preincubation with both hEphB3-Fc and hPlxdc2-FcStrep further reduced RRV-YFP wt infection on HaCaT cells, when compared to preincubation with either hEphB3-Fc or hPlxdc2-FcStrep alone (Fig 2C). While preincubation with EphB3-Fc did not reduce RRV-YFP gH-AELAAN infection, preincubation with either hPlxdc2-FcStrep or a combination of hPlxdc2-FcStrep and hEphB3-Fc reduced infection by approx. 50% as observed for RRV-YFP wt infection (Fig 2C). Infection of SLK cells and rhesus monkey fibroblasts was also slightly decreased by preincubation of the viral inoculum with hPlxdc2-FcStrep, infection of SLK by RRV-YFP wt to 63.5% ± 3.7% and infection of RF by RRV-YFP wt to 73.9% ± 11.3% relative to preincubation with FcStrep as control. However, this effect was less pronounced than on HaCaT and less pronounced than the effect of hPlxdc2-FcStrep on RRV-YFP gH-AELAAN infection of the same cell types (Fig 2D). Taken together, the immunoprecipitation and blocking experiments confirm the independence of the gH-Plxdc interaction of the previously described Eph interaction motif (23) and EphB3-binding.

**Figure 2.**
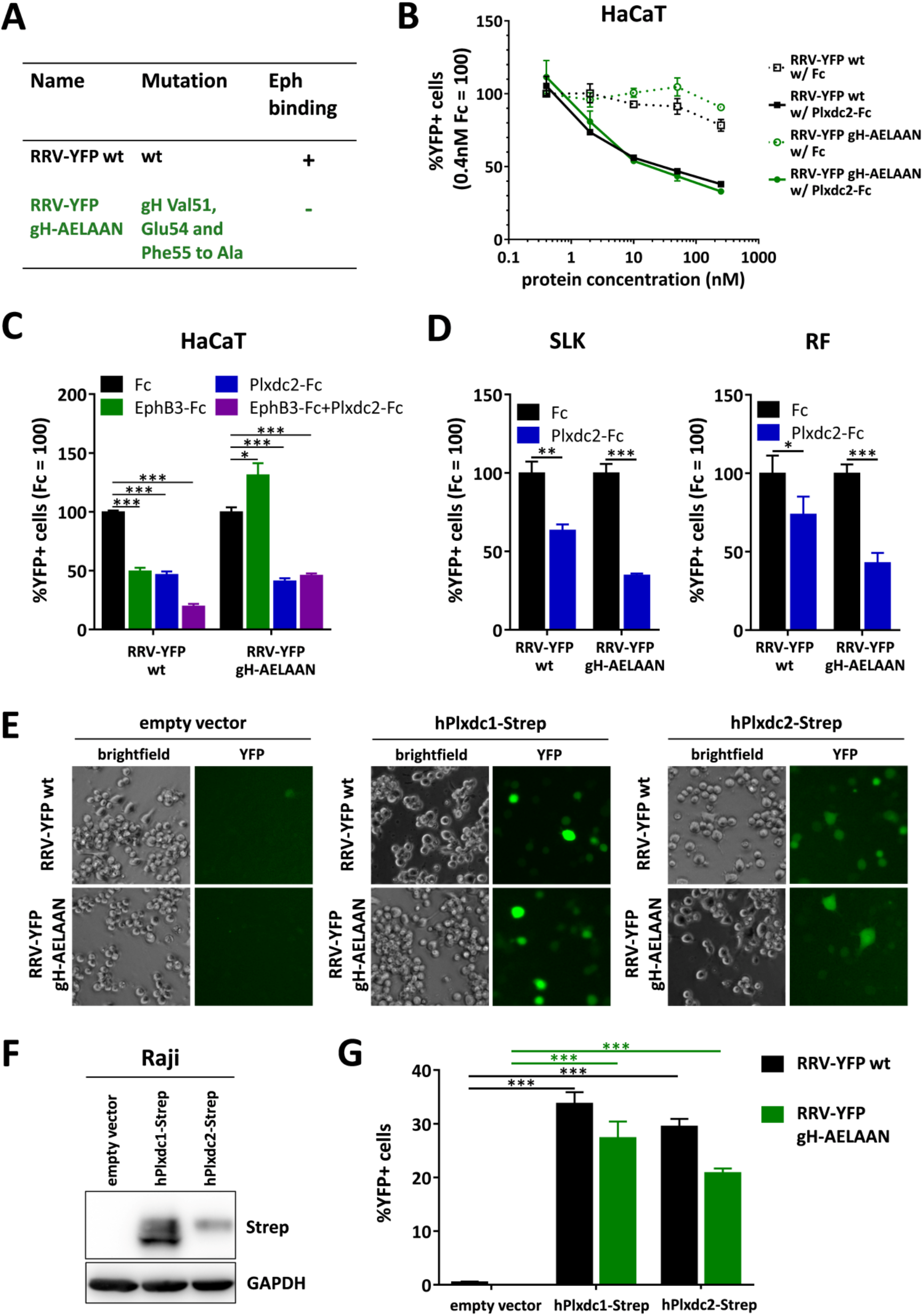
Plxdc1/2 function as entry receptors for RRV. **A)** List of BAC-derived recombinant viruses and introduced mutations used in this figure. **B)** Dose-dependent inhibition of RRV 26-95 infection by soluble human Plxdc2-FcStrep on HaCaT cells. RRV-YFP wt and RRV-YFP gH-AELAAN were pre-incubated with hPlxdc2-FcStrep for 30min at room temperature. FcStrep alone was used as control. YFP expression as indicator of infection was measured by flow cytometry. Infection in the presence of 0.4nM FcStrep was set to 100% (MOI ~0.2, triplicates, error bars represent SD). **C)** Inhibition of RRV 26-95 infection by soluble Plxdc2-FcStrep and EphB3-Fc on HaCaT cells. RRV-YFP wt and RRV-YFP gH-AELAAN were pre-incubated with 100nM hPlxdc2-FcStrep, 10nM EphB3-Fc or a combination of 100nM hPlxdc2-FcStrep and 10nM EphB3-Fc for 30min at room temperature. FcStrep alone was used as control. YFP expression as indicator of infection was measured by flow cytometry. Infection with FcStrep was set to 100% (MOI ~0.2, triplicates, error bars represent SD). **D)** Inhibition of RRV 26-95 infection by soluble human Plxdc2-FcStrep on SLK cells and rhesus fibroblasts. RRV-YFP wt and RRV-YFP gH-AELAAN were pre-incubated with 250nM hPlxdc2-FcStrep for 30min at room temperature. FcStrep alone was used as control. YFP expression as indicator of infection was measured by flow cytometry. Infection with FcStrep was set to 100% (MOI ~0.05-0.1, triplicates, error bars represent SD). **E)** Raji cells were transduced with TwinStrep-tagged human Plxdc1 and Plxdc2 (hPlxdc1-Strep/ hPlxdc2-Strep) expression constructs or an empty vector control, briefly selected and infected with RRV-YFP wt or RRV-YFP gH-AELAAN normalized to genome copies as determined by qPCR. Micrographs show representative infection of the indicated cell pools. **F)** Lysates of transduced Raji cell pools were analyzed for Plxdc1/2-Strep expression by Western blot. **G)** Quantification of (E) by flow cytometric analyses of YFP reporter gene expression as indicator of infection.

To establish receptor function, we performed gain-of-function experiments using ectopic Plxdc1/2 overexpression. Raji cells were transduced with lentiviruses encoding TwinStrep-tagged human Plxdc1/2 constructs (hPlxdc1-Strep/ hPlxdc2-Strep). The EBV-positive, human lymphoblast cell line only allows for low-level RRV 26-95 infection even with amounts of input virus corresponding to high MOI on adherent cells like SLK, HaCaT or RF. Therefore, changes in susceptibility to infection mediated by Plxdc1/2 overexpression should be readily detectable and allow for a clear differentiation of the contribution of Plxdc1/2 to RRV infection over the very low intrinsic susceptibility to infection. Indeed, ectopic expression of hPlxdc1/2-Strep increased RRV-YFP wt and RRV-YFP gH-AELAAN infection 40 to 60-fold from approx. 0.5% basal infection dependent on the expression plasmid and RRV strain (Fig 2E, G). However, we did not observe pronounced differences in effects mediated by hPlxdc1-Strep or hPlxdc2-Strep, indicating no clear Plxdc receptor preference of RRV 26-95.

In a next step we characterized the Plxdc binding motif on RRV gH. As the RRV 17577 gH-Plxdc2 interaction depends on gL whereas the RRV 26-95 gH-Plxdc2 interaction does not, we focused on the N-terminal domain I of gH which, in analogy to EBV gH/gL (34), most likely constitutes the gL-binding interface. The differences in the Plxdc interaction of RRV isolates 26-95 and 17577 as well as the lack of an interaction between KSHV gH/gL and Plxdc receptors suggested a motif that is only partially conserved between the RRV isolates and missing in KSHV. Using sequence comparisons (Fig 3A) we identified a putative interaction motif spanning 7 or 6 amino acid motif in the N-terminal region of RRV 26-95 gH and RRV 17577 gH, respectively, that is not conserved in KSHV gH. The motif is located close to the Eph-interaction motif we described previously, facing in the opposite direction in a homology model of the RRV 26-95 gH/gL complex based on the EBV gH/gL crystal structure (3PHF) (Fig 3B). Deletion of this motif completely abrogated the gH/gL interaction with Plxdcs of both RRV 26-95 (Fig 3C) and RRV 17577 (Fig 3D). To further characterize the contribution of individual residues in the ‘Tyr(Y)-Glu(E)-Tyr(Y)-Asn(N)-Glu(E)-Glu(E)-Lys(K)’ (RRV 26-95) motif we performed single amino acid substitutions to alanine. The ability of mutant RRV 26-95 gH-V5 to bind myc-tagged Plxdc1/2 of human (hPlxdc1/2-myc) (Fig 3E) or rhesus macaque origin (mmPlxdc1/2-myc) (Fig 3F) was analyzed by immunoprecipitation of gH via the V5-tag and Western blot. While several single amino acid substitutions decreased the interaction of RRV gH with Plxdcs to some degree, residues Tyr23 and Glu25 that are conserved in isolates 26-95 and 17577 appear to be critical for the interaction of gH with human and rhesus macaque Plxdc1 and Plxdc2. Furthermore, substitution of glutamate with alanine at position 22, which is not conserved between isolates 26-95 and 17577, had a pronounced, albeit slightly weaker effect on the gH-Plxdc1/2 interaction (Fig 3E, F).

**Figure 3.**
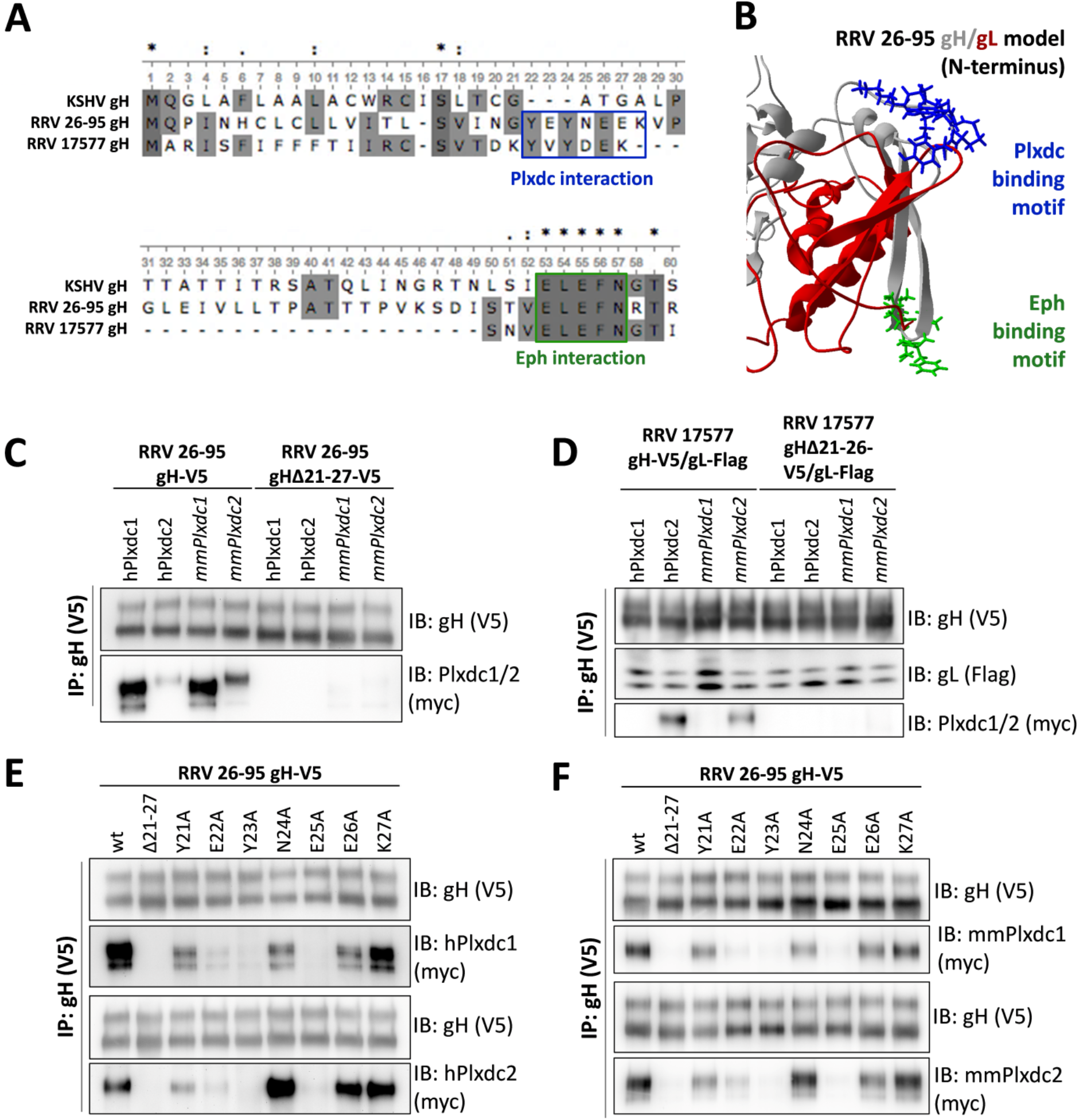
An amino acid sequence motif in the N-terminal region of RRV 26-95 and RRV 17577 gH is essential for the Plxdc interaction. **A)** Multiple sequence alignment of the N-terminal region of gH of KSHV and the two RRV isolates 26-95 and 17577. Boxes indicate the binding motives for Eph receptors (green) and Plxdc receptors (blue). **B)** Homology-based structure prediction of the RRV 26-95 gH/gL complex based on the crystal structure of the EBV gH/gL complex (PDB number 3PHF) using the Iterative Threading ASSembly Refinement (I-TASSER) server and the CO-THreader (COTH) algorithms for protein-protein complex structure and multi-chain protein threading. The Eph receptor interaction motif is shown in green, the Plxdc interaction motif is shown in blue, gL is shown in red, gH is shown in grey. **C)** Deletion of the Plxdc interaction motif in RRV 26-95 gH (amino acid 21-27, “YEYNEEK”) abrogates gH interaction with Plxdc1 and Plxdc2. V5-tagged gH wt or gHΔ21-27 were immunoprecipitated in the presence of full-length human or macaca mulatta Plxdc1-myc or Plxdc2-myc using monoclonal antibody to the V5-tag. Precipitates were analyzed by Western blot. **D)** Deletion of the Plxdc interaction motif in RRV 17577 gH (amino acid 21-26, “YVYDEK”) abrogates gH interaction with Plxdc2. V5-tagged gH wt or gHΔ21-26 was co-expressed with Flag-tagged RRV 17577 gL. gH-V5/gL-Flag complexes were immunoprecipitated in the presence of full-length human or macaca mulatta Plxdc1-myc or Plxdc2-myc using monoclonal antibody to the V5-tag. Precipitates were analyzed by Western blot. **E)** Mutational scan of Plxdc interaction motif (amino acid 21-27, “YEYNEEK”) of RRV 26-95 gH identifies human Plxdc1/2-interacting residues. V5-tagged gH mutants were immunoprecipitated in the presence of full-length human Plxdc1-myc or Plxdc2-myc using monoclonal antibody to the V5-tag. Precipitates were analyzed by Western blot. RRV gHΔ21-27 serves as negative control. **F)** Mutational scan of Plxdc interaction motif (amino acid 21-27, “YEYNEEK”) of RRV 26-95 gH identifies rhesus macaque Plxdc1/2-interacting residues. V5-tagged gH mutants were immunoprecipitated in the presence of full-length human Plxdc1-myc or Plxdc2-myc using monoclonal antibody to the V5-tag. Precipitates were analyzed by Western blot. RRV gHΔ21-27 serves as negative control. Abbreviations: IP: immunoprecipitation, IB: immunoblotting, h: human, mm: macaca mulatta (rhesus macaque).

To further analyze the contribution of the gH/gL-Plxdc interaction in the context of infection we constructed virus mutants deleted in the seven amino acid interaction motif in the background of RRV-YFP 26-95 wildtype (RRV-YFP gHΔ21-27), and in the background of an RRV-YFP 26-95 strain mutated in the Eph-interaction motif described previously by our group (RRV-YFP gH-AELAAN, RRV-YFP gHΔ21-27-AELAAN) using a two-step, lambda red-mediated recombination system (35) (Fig 4A). Blocking experiments using soluble hPlxdc2-FcStrep decoy receptor on HaCaT cells confirmed that deletion of the seven amino acid motif was sufficient to abrogate the gH-Plxdc2 interaction on viral particles (Fig 4B). While infection of RRV-YFP wt and RRV-YFP gH-AELAAN was inhibited by approx. 60% to 70% respectively, infection of RRV-YFP gHΔ21-27 and RRV-YFP gHΔ21-27-AELAAN was not affected even by high concentrations of soluble hPlxdc2-FcStrep (Fig 4B). All infections were carried out at approx. MOI 0.05. Analogously, the Eph- or Plxdc-receptor-binding-negative RRV mutants were no longer inhibited by preincubation with the respective soluble receptor (hEphB3-Fc or hPlxdc2-FcStrep) in single or double inhibition experiments on HaCaT cells (Fig 4C). The receptor-specificity conveyed by the respective interaction motif was further analyzed in lentiviral vector-mediated Plxdc1/2-Strep overexpression experiments in Raji B lymphocytes. EphA7, which had previously been described by our group to be critical for RRV infection of BJAB B lymphocytes, was used as control for Eph-mediated infection. Expression of Plxdc1/2-Strep as well as EphA7-Strep dramatically enhanced susceptibility of Raji cells (Fig 4D, E). RRV-YFP wt infection increased from 0.14 ± 0.04% on empty vector transduced Raji cells to approx. 8.5% upon EphA7 overexpression and to approx. 17% upon Plxdc1/2 overexpression, without pronounced differences between the Plxdc family members. Mutation of the Eph-interaction motif in RRV-YFP gH-AELAAN completely abrogated the gain in susceptibility on EphA7 overexpressing cells while mutation of the Plxdc-interaction motif completely abrogated the gain in susceptibility on Plxdc1/2 overexpressing cells, confirming selective knockout of each individual receptor interaction in the respective mutant. Furthermore, deletion of the Eph-interaction motif in RRV-YFP gH-AELAAN did not impact the infection of Plxdc1/2 overexpressing cells in comparison to RRV-YFP wt infection. In contrast, we observed an approx. 2-fold higher infection of RRV-YFP gHΔ21-27 on EphA7 overexpression cells, when compared to RRV-YFP wt. Together, the blocking and overexpression experiments indicate an independent rather than cooperative nature of Eph and Plxdc receptor function.

**Figure 4.**
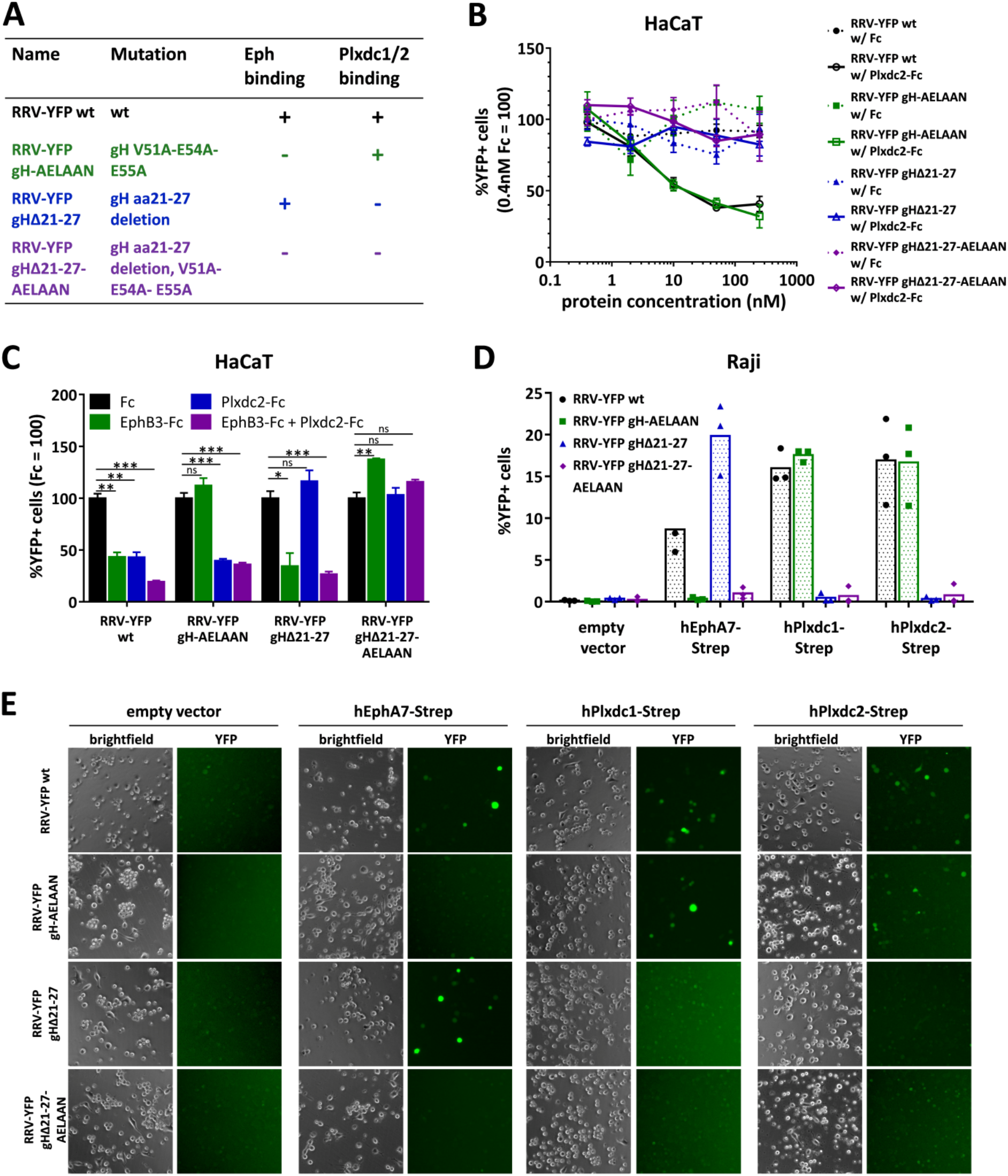
Deletion of the seven amino acid Plxdc-binding motif is sufficient to detarget RRV 26-95 from Plxdc receptors. **A)** List of BAC-derived recombinant viruses and introduced mutations used in this figure. **B)** Dose-dependent inhibition of RRV 26-95 infection by soluble human Plxdc2-FcStrep on HaCaT cells. RRV-YFP wt, RRV-YFP gH-AELAAN, RRV-YFP gHΔ21-27 and RRV-YFP gHΔ21-27-AELAAN were pre-incubated with hPlxdc2-FcStrep for 30min at room temperature. FcStrep alone was used as control. YFP expression as indicator of infection was measured by flow cytometry. Infection in the presence of 0.4nM FcStrep was set to 100% (MOI ~0.05, triplicates, error bars represent SD). **C)** Inhibition of RRV 26-95 infection by soluble Plxdc2-FcStrep and EphB3-Fc on HaCaT cells. RRV-YFP wt, RRV-YFP gH-AELAAN, RRV-YFPgHΔ21-27 and RRV-YFP gHΔ21-27-AELAAN were pre-incubated with 100nM hPlxdc2-FcStrep, 10nM EphB3-Fc or a combination of 100nM hPlxdc2-FcStrep and 10nM EphB3-Fc for 30min at room temperature. FcStrep alone was used as control. YFP expression as indicator of infection was measured by flow cytometry. Infection with FcStrep was set to 100% (MOI ~0.1-0.2, triplicates, error bars represent SD). **D)** Raji cells were transduced with TwinStrep-tagged human EphA7, Plxdc1 or Plxdc2 (hEphA7-Strep, hPlxdc1-Strep, hPlxdc2-Strep) expression constructs or an empty vector control, briefly selected and infected with RRV-YFP wt, RRV-YFP gH-AELAAN, RRV-YFP gHΔ21-27 or RRV-YFP gHΔ21-27-AELAAN normalized to genome copies as determined by qPCR. YFP expression as indicator of infection was measured by flow cytometry. The mean across three independent sets of RRV stocks is indicated by columns. The means of individual triplicate infections for each set of RRV stocks are given as symbols within the respective columns. **E)** Micrographs show representative infection of one set of RRV stocks in (D).

To quantitatively analyze the contribution of the Plxdc1/2-interaction to RRV infection of different cell types, RRV-YFP wt and RRV-YFP receptor binding mutant inocula were normalized to genome copies as determined by qPCR, and target cells were inoculated with the same number of encapsidated input virus genomes for wt and each mutant virus strain in an MOI range of 0.05 to 1 on adherent cells. Infection as determined by the percentage of YFP+ cells was normalized to RRV-YFP wt, which was set to 1. For suspension cell lines, experiments with RRV-YFP wt infection over 1% were included in the analysis. Four (three for RF, five for MFB5487) independent sets of RRV wt and mutant stocks were used to compensate for variability in stock preparation. None of the analyzed adherent cell lines showed a preferential use of Plxdc receptors over Eph receptors based on the reduction of specific infectivity of Eph-binding and Plxdc-interaction-deficient mutants (Fig 5A). On rhesus monkey fibroblasts (RF), compared to RRV-YFP wt, RRV-YFP gHΔ21-27, RRV-YFP gH-AELAAN and RRV-YFP gHΔ21-27-AELAAN exhibited a decrease in infection of approx. 30%, 50% and 70%, respectively. Similarly on HaCaT, infection with RRV-YFP gH-AELAAN and RRV-YFP gHΔ21-27-AELAAN was reduced by approx. 60% and 75%, respectively, compared to RRV-YFP wt. While RRV-YFP gHΔ21-27 exhibited a defect, comparable to RRV-YFP gH-AELAAN in three of the analyzed independent sets of RRV stocks, this reduction in infectivity did not reach significance due to one outlier. On SLK cells, mutation of the Eph-interaction motif led to an approx. 65% decrease in infection, while RRV-YFP gHΔ21-27 infection was on average comparable to RRV-YFP wt infection. In contrast, we identified B lymphocyte lines of human and macaque origin that exhibit a preference in the receptor usage for either Eph or Plxdc family members (Fig 5B). When normalized to genome copies, RRV-YFP gHΔ21-27 infection on human Raji lymphoblasts was comparable to RRV wt infection while mutation of the Eph-interaction motif lead to an approx. 90% decrease in infection, indicating preferential infection through Eph family receptors, albeit at low levels, as mentioned above. Similarly, RRV-YFP gHΔ21-27 exhibited only a minor defect (approx. 20% reduced infection) on immortalized B lymphocytes of macaca mulatta origin (MMB1845) that did not reach significance, while infection with RRV-YFP gH-AELAAN and gHΔ21-27-AELAAN was reduced by approx. 90% in comparison to RRV-YFP wt, again indicating preferential infection through Eph family receptors. Conversely, deletion of the Plxdc-interaction motif decreased infection of a B lymphocyte cell line of macaca fascicularis origin (MFB5487) by approx. 75% whereas RRV-YFP gH-AELAAN exhibited an approx. 50% defect, indicating preferential use of the Plxdc interaction for infection of MFB5487 by RRV. Mutation of both the Eph- and Plxdc-interaction motif led to an even more pronounced defect of approx. 90%.

**Figure 5.**
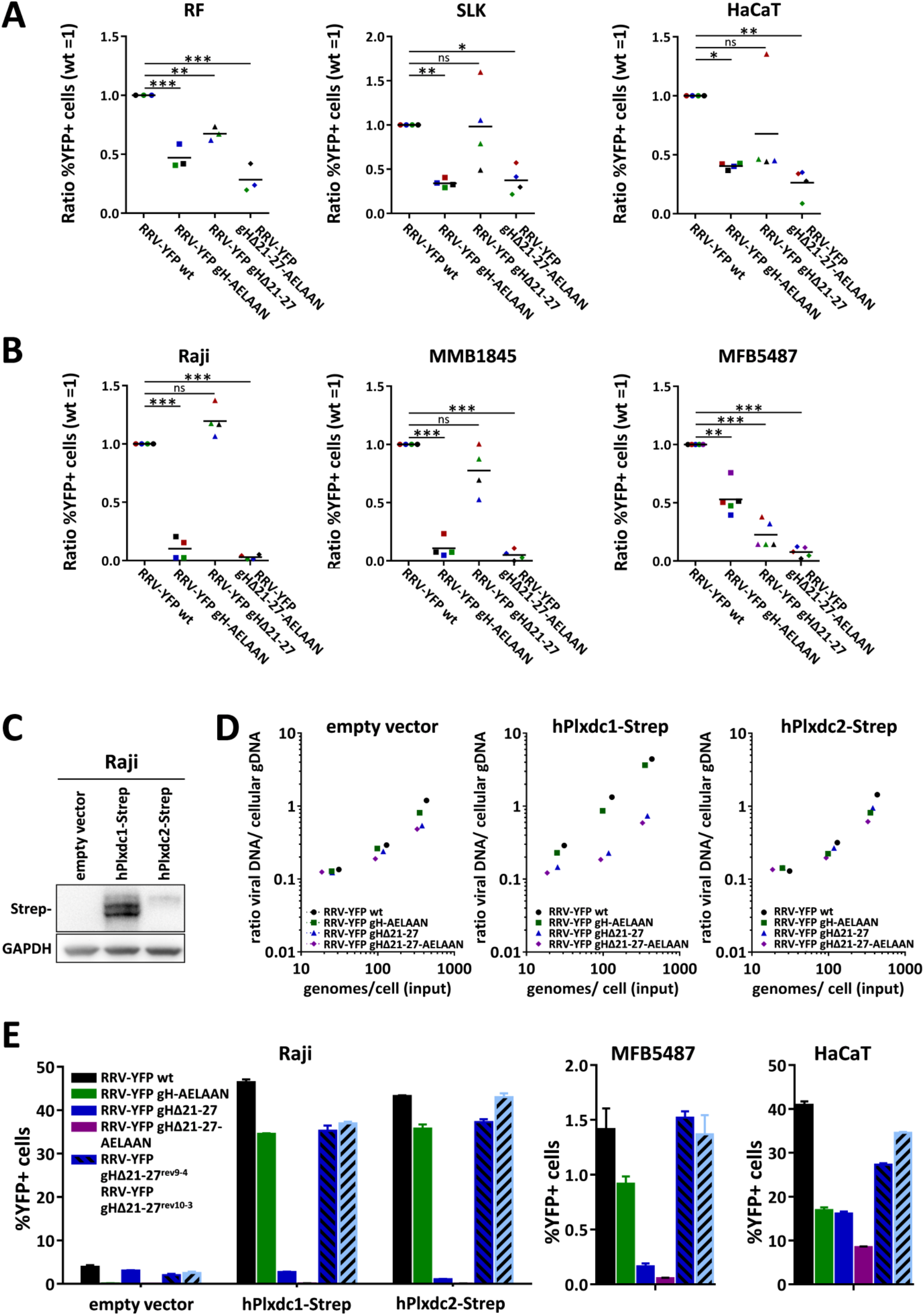
The contribution of the RRV 26-95 gH-Plxdc interaction to infection is cell type-specific and may in part be dependent on attachment effects. **A)** RRV 26-95 deleted in the Plxdc interaction motif exhibits reduced specific infectivity on HaCaT and RF, but not SLK cells. Target cells were infected with RRV-YFP wt, RRV-YFP gH-AELAAN, RRV-YFP gHΔ21-27 or RRV-YFP gHΔ21-27-AELAAN normalized to genome copies as determined by qPCR. YFP expression as indicator of infection was measured by flow cytometry and normalized to RRV-YFP wt infection. Means of individual normalized infections with three (RF) or four (HaCaT, SLK) independent sets of RRV stocks are given as symbols of different color. Sets with RRV-YFP wt infection in an MOI range of 0.05 - 1 were used for analysis. The mean across the independent sets of RRV stocks is indicated by black lines. **B)** RRV 26-95 deleted in the Plxdc interaction motif exhibits reduced specific infectivity on MFB5487, but not Raji and MMB1845 cells. Target cells were infected with RRV-YFP wt, RRV-YFP gH-AELAAN, RRV-YFP gHΔ21-27 or RRV-YFP gHΔ21-27-AELAAN normalized to genome copies as determined by qPCR. YFP expression as indicator of infection was measured by flow cytometry and normalized to RRV-YFP wt infection. Means of individual normalized infections with four (Raji, MMB1845) or five (MFB5487) independent sets of RRV stocks are given as symbols of different color. Sets with RRV-YFP wt infection exceeding 1% were used for analysis. The maximal achieved RRV-YFP wt infection was 6.1% for Raji, 5.6% for MMB1845 and 3.1% for MFB5487, respectively. The mean across the independent sets of RRV stocks is indicated by black lines. **C)** Western blot analysis of Raji cells transduced with TwinStrep-tagged human Plxdc1 and Plxdc2 (hPlxdc1-Strep/ hPlxdc2-Strep) expression constructs or an empty vector control. **D)** Attachment of RRV 26-95 on transduced Raji cells is affected by hPlxdc1-Strep, but not hPlxdc2-Strep overexpression. Cells, analyzed in (C), were incubated with cold virus at the indicated concentrations at 4°C for 30min followed by genomic DNA isolation. The ratio of viral to cellular DNA as measurement for attached virus was calculated based on ΔCt values of a genomic (CCR5) and a viral locus (ORF73/ LANA) as determined by qPCR and plotted against input viral genome number. **E)** Re-introduction of the seven amino acid motif crucial for Plxdc interaction rescues RRV-YFPgHΔ21-27 infection. Transduced Raji cells, analyzed in (C), MFB5487 and HaCaT cells were infected with RRV-YFP wt, RRV-YFP gH-AELAAN, RRV-YFP gHΔ21-27, RRV-YFP gHΔ21-27-AELAAN or two RRV-YFP gHΔ21-27 revertants (RRV-YFP gHΔ21-27^rev9-4^, RRV-YFPgHΔ21-27^rev10-3^) normalized to genome copies as determined by qPCR. YFP expression as indicator of infection was measured by flow cytometry (triplicates, error bars represent SD).

To evaluate the contribution of potential attachment effects on the observed differences in specific infectivity, we analyzed the capacity of virions to bind Plxdc1/2 overexpressing Raji cells in comparison to empty vector transduced Raji cells (Fig 5C, D). It should be noted that transduction with an Plxdc1 encoding lentiviral vector resulted in drastically higher levels of protein expression as assessed by Western blot analysis than transduction with an Plxdc2 encoding lentiviral vector (Fig 5C). The ratio of cell-bound viral DNA to input genomes was used as a surrogate marker for virus attachment. Mutation of the Plxdc- and Eph-interaction motif had no detectable effect on attachment to control vector-transduced and Plxdc2-overexpressing Raji cells. In contrast, Raji cells overexpressing Plxdc1 showed increased attachment of RRV-YFP wt and RRV-YFP gH-AELAAN, while attachment of Plxdc-interaction negative RRV-YFPmutants gHΔ21-27 and gHΔ21-27-AELAAN was not enhanced and remained comparable to empty vector control and Plxdc2-overexpressing cells.

To exclude effects of potential offsite genomic rearrangements in RRV-YFP gHΔ21-27 we created two independent revertants (RRV-YFP gHΔ21-27^rev9-4^ and RRV-YFP gHΔ21-27^rev10-3^). Restoration of the residues deleted in RRV-YFP gHΔ21-27 restored infection on hPlxdc1/2-transduced Raji, MFB5487 and HaCaT cells to RRV-YFP wt levels, with no pronounced differences between RRV-YFP wt, RRV-YFP gHΔ21-27^rev9-4^ and RRV-YFP gHΔ21-27^rev10-3^ (Fig 5E). Similar to the approx. 11-fold increase of RRV-YFP wt infection upon hPlxdc1/2 overexpression, infection with RRV-YFP gHΔ21-27 revertants increased from 2.00% ± 0.30% (2.54% ± 0.29%) on empty vector transduced Raji cells to 35.25% ± 1.21% (36.99% ± 0.39%) and 37.27% ± 0.68% (43.00% ± 0.99%) upon hPlxdc1 and hPlxdc2 overexpression for RRV-YFP gHΔ21-27^rev9-4^ (RRV-YFP gHΔ21-27^rev10-3^), respectively (Fig 5E).

Our interaction data (Fig 1A, B) suggests that at least RRV 26-95 gH can bind Plxdc1/2 in the absence of gL. We therefore aimed to analyze if infection with a RRV 26-95 gL deletion mutant (RRV-YFP ΔgL) is influenced by Plxdc1 and 2 expression to a similar degree as RRV-YFP wt and RRV-YFP gH-AELAAN infection. As shown before, overexpression of hEphA7-Strep or hPlxdc1/2-Strep in Raji cells dramatically increased RRV-YFP wt infection compared to empty vector transduced cells (22-fold for hEphA7-Strep, 23-fold for hPlxdc2-Strep and 26-fold for hPlxdc2-Strep) (Fig 6). RRV-YFP gH-AELAAN infection was increased from 0.25% ± 0.05% basal infection on empty vector transduced cells to 52.53% ± 2.06% and 65.5% ± 1.22%, respectively, upon hPlxdc1-Strep or hPlxdc2-Strep overexpression, while hEphA7-Strep overexpression had no pronounced effect on infection. We observed a pattern similar to RRV-YFP gH-AELAAN for the infection with two independent RRV-YFP ΔgL clones, RRV-YFP ΔgL^3-3^ and RRV-YFP ΔgL^3-5^. While, hEphA7-Strep overexpression did not influence RRV-YFP ΔgL infection, hPlxdc1/2-Strep overexpression enhanced infection from 0.36% ± 0.03% (0.48% ± 0.04%) on empty vector transduced Raji cells to 31.05% ± 0.95% (36.25% ± 1.42%) and 35.59% ± 0.94% (42.55% ± 0.57%) upon hPlxdc1 and hPlxdc2 overexpression for RRV-YFP ΔgL^3-3^ (RRV-YFP ΔgL^3-5^) respectively (Fig 6C). Even though the effect of Plxdc1/2 overexpression on RRV-YFP ΔgL infection was slightly less pronounced than the effect on RRV-YFP wt and RRV-YFP gH-AELAAN infection, this data demonstrates that the use of Plxdc1 and Plxdc2 as RRV 26-95 entry receptors is possible in a gL-independent manner.

**Figure 6.**
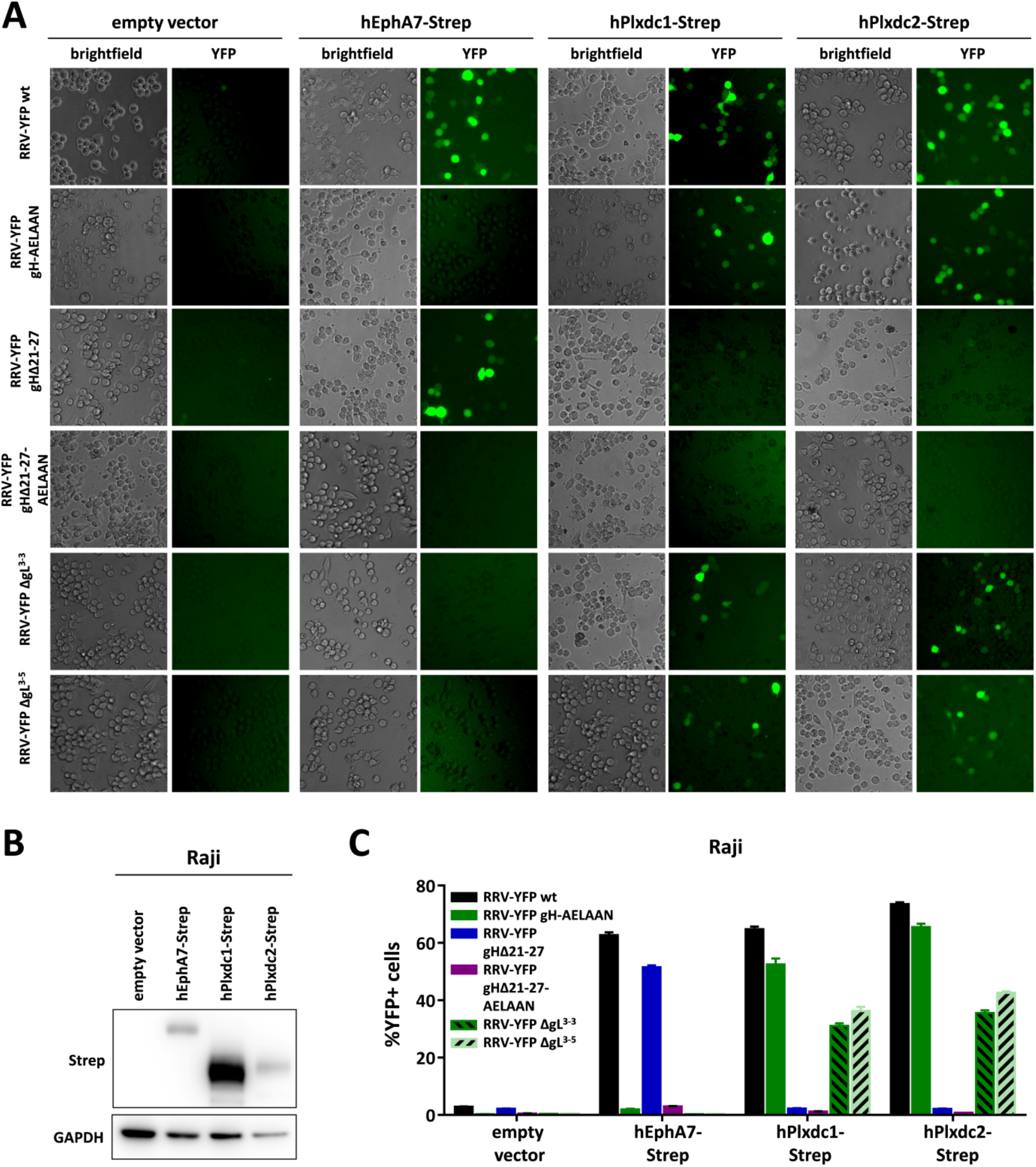
Plxdc1/2-dependent infection with RRV 26-95 does not require gL. **A)** Raji cells were transduced with TwinStrep-tagged human EphA7, Plxdc1 and Plxdc2 (hEphA7-Strep/ hPlxdc1-Strep/ hPlxdc2-Strep) expression constructs or an empty vector control, briefly selected and infected with RRV-YFP wt or RRV-YFP gH-AELAAN, RRV-YFP gHΔ21-27 or one of two RRV-YFP ΔgL clones normalized to genome copies as determined by qPCR. Micrographs show representative infection of the indicated cell pools. **B)** Lysates of transduced Raji cell pools were analyzed for EphA7-Strep and Plxdc1/2-Strep expression by Western blot. **C)** Quantification of (A) by flow cytometric analyses of YFP reporter gene expression as indicator of infection (triplicates, error bars represent SD).

## DISCUSSION

In this study we identified the Plexin domain containing proteins 1 and 2 as a novel family of entry receptors for RRV. Plxdc1/2 interact with the gH/gL complex of RRV in a region close to the previously characterized binding motif for Eph receptors. While the Eph-interaction as well as the critical Eph binding motif in domain I of gH is conserved between RRV and the closely related human pathogenic KSHV (23), the interaction with Plxdc1/2 is exclusive to RRV and even exhibits differences between isolates 26-95 and 17577 as prototypic members of the two described RRV sequence clades.

According to our results, differences between Plxdc1 and Plxdc2 may also exist in terms of function. While overexpression of both Plxdc1 and Plxdc2 in Raji cells, that were virtually “non-susceptible” under the conditions used, lead to robust RRV 26-95 infection (Fig 4D-E), only overexpression of Plxdc1 enhanced attachment of RRV wt and RRV gH-AELAAN in comparison to either non-transduced Raji cells or Plxdc-binding deficient RRV mutants (Fig 5D). Whether this is primarily due to differences in expression levels, consistently observed between Plxdc1 and Plxdc2 upon lentiviral overexpression or due to underlying functional differences in the gH interaction with these molecules remains to be determined.

Mutation of residues Tyr23 and Glu25 in gH domain I, which are conserved between RRV isolate 26-95 and 17577, almost abolished the interaction with Plxdc2, which is bound by gH/gL of both isolates. Although the same residues are critical for the interaction of RRV gH 26-95 with Plxdc1, the partial conservation of the region appears to be insufficient to confer binding of 17577 gH to Plxdc1. Interestingly, the interaction of 17577 gH with Plxdc2 is dependent on the presence of gL in the gH/gL complex, whereas 26-95 gH is able to bind Plxdc1/2 independent of gL. These features of Plxdc1/2 binding specificities appear similar to the interaction of the gH/gL complex with Eph receptors, wherein mutation of the strictly conserved Eph interaction motif is sufficient to abrogate receptor binding, but differences in KSHV and RRV affinities for A- and B-type Eph RTKs indicate the existence of additional regions in gH or gL that contribute to or modulate the interaction. Whether these preferences for different members of conserved receptor families also influence e.g. cell or tissue tropism, viral spread or pathogenicity has not been determined. Sequence comparisons of over twenty RRV isolates identified dramatic differences in the extracellular domains of gH as well as in gL between isolates that fell in two discrete groupings either similar to 26-95 or 17577, while variation in other glycoproteins R1, gM, gN, orf68 was minimal between clades (4). However, if these clade-specific glycoprotein variations influence the observed differences in pathogenicity between RRV strains 26-95 and 17577 remains to be seen. Another interesting question would be if the receptor-binding function of the N-terminal region of gH is conserved between RRV and KSHV, and probably EBV, and whether KSHV and EBV can bind another receptor through this region. So far, we did not identify corresponding interactions for KSHV, but in principle the region of gH encompassing the Plxdc2 binding site is present in both viruses and it is tempting to assume some functional conservation.

Along those lines, the evolutionary factors that drive interaction with different receptor families and the resulting multitude of herpesvirus–receptor interactions is highly interesting. For e.g. EBV and HCMV, a clear correlation between receptor usage, dependent on viral interaction partners of the gH/gL complex, and cell tropism has been demonstrated in various studies ((36), reviewed in (37)). However, for rhadinoviruses, the picture is less clear. We demonstrate that RRV infection of select cell lines exhibited a dependence on specific receptors, e.g. Raji infection – to the modest extent that was possible without recombinant receptor overexpression – was dependent on the gH/gL-Eph interaction, MFB5487 infection was more dependent on the Plxdc-interaction (Fig 5), and similarly mutation of the Eph-interaction motif did not impact infection on all cell types equivalently (23). Yet, a definite correlation between exclusive receptor usage and infection of specific cell types is not obviously apparent. While the notion of a role of different receptor interactions in KSHV and RRV cell and tissue tropism is tempting and an established concept for related viruses (37–39), the possibility of a redundant function should not be discarded. Redundancy could be driven by the need to escape antibodies e.g. to one receptor binding site. For instance, *in vitro* infection of human keratinocytes or rhesus fibroblasts seemed to be impacted to a similar degree by either deletion of the Eph- or of the Plxdc-interaction motif (Fig 5). To ultimately address the correlation between receptor- and tissue-tropism, *in vivo* studies using receptor-de-targeted mutants to analyze cell and tissue tropisms will be required. The importance of *in vivo* studies is also supported by a recent report that showed that a gL-null RRV mutant still established persistent infection in the B cell compartment upon intravenous inoculation, while infection of B cells *in vitro* was drastically reduced (24), a finding that could be explained by the gL-independent usage of Plxdc1/2 for B cell infection *in vivo.* At a minimum, our data on RRV-YFP ΔgL infection of Plxdc1 and Plxdc2 overexpressing Raji cells (Fig 6) confirms that RRV can efficiently use Plxdc1 and Plxdc2 as entry receptors in the absence of gL. Finally, the fact that a mutant that was deleted in both the Eph and the Plxdc1/2 interaction motif, RRV-YFP gHΔ21-27-AELAAN, was still able to replicate on rhesus monkey fibroblasts, as evidenced by the fact that we were able to grow a virus stock on these cells, and was still infectious to a certain degree on a number of cell lines (Figs 4 and 5) indicates a surprisingly high degree of redundancy in the entry pathways that RRV can use and hints at the existence of additional receptors or host factors besides Ephs and Plexin domain containing proteins.

On a more speculative note, the apparent overlap between virus receptors and tumor-associated membrane proteins may represent an interesting research subject. Eph receptors were first identified in an attempt to characterize tyrosine kinases involved in cancer (40) and altered expression in various cancer types has been demonstrated for several Eph family members (reviewed in (41)). Similarly, Plxdc1 was first described in a screen for novel tumor endothelial members (25) and expression of both Plxdc1 and Plxdc2 is elevated in the endothelium of solid tumors (25, 26, 42, 43). Plxdc1 expression has been described as prognostic marker and modulating factor for various human cancers (42–46). Given the overlap between the required changes e.g. in metabolism, transcription, and signaling for cancer growth and virus replication it seems not unlikely that either elevated expression or signaling of these molecules is favorable for both virus infection and cancer progression.

## MATERIAL AND METHODS

### Cells

Human embryonic kidney (HEK) 293T cells (RRID:CVCL_0063) (laboratory of Tobias Moser), SLK cells (RRID:CVCL_9569) (NIH AIDS Research and Reference Reagent program), rhesus monkey fibroblasts (RF) (laboratory of Prof. Rüdiger Behr) and HaCaT human keratinocytes (RRID:CVCL_0038) were cultured in Dulbecco’s Modified Eagle Medium (DMEM), high glucose, GlutaMAX, 25mM HEPES (Thermo Fisher Scientific) supplemented with 10% fetal calf serum (FCS) (Thermo Fisher Scientific), and 50μg/ml gentamycin (PAN Biotech). iSLK cells (laboratory of Don Ganem, Novartis Institutes for BioMedical Research, Emeryville, CA, USA) were maintained in DMEM supplemented with 10% FCS, 50μg/ml gentamycin, 2.5μg/ml puromycin (InvivoGen) and 250μg/ml G418 (Carl Roth). Raji cells (RRID:CVCL_0511) (laboratory of Jens Gruber), MFB5487 (a clonal cell line established from macaca fascicularis PBMC, immortalized by infection with herpesvirus papio; a kind gift from Ulrike Sauermann) and MMB1845 cells (a clonal cell line established from macaca mulatta PBMC, immortalized by infection with herpesvirus papio; a kind gift from Ulrike Sauermann) were cultured in RPMI (Thermo Fisher Scientific) supplemented with 10% FCS and 50μg/ml gentamycin.

### BAC mutagenesis and virus production

RRV recombinants (RRV-YFP gHΔ21-27, RRV-YFP ΔgL and RRV-YFP gHΔ21-27-AELAAN) were generated based on BAC35-8 (47) and RRV gH-AELAAN (23) respectively, using a two-step, markerless λ-red-mediated BAC recombination strategy as described by Tischer et al. (35). RRV-YFP ΔgL harbors a 128 bp deletion, which introduces a frameshift after amino acid 26 and a stop codon after amino acid 37, leaving only six amino acids of the original gL sequence after the putative signal peptide cleavage site. RRV-YFP gHΔ21-27 revertants were generated based on RRV-YFP gHΔ21-27 following the same protocol described by Tischer et al. (35). In short, recombination cassettes were generated from the pEPKanS template by polymerase chain reaction (PCR) with Phusion High Fidelity DNA polymerase (Thermo Fisher Scientific) using long oligonucleotides (Ultramers; purchased from Integrated DNA Technologies (IDT)) (see S1 Table for a complete list of primers). Recombination cassettes were transformed into RRV-YFP-carrying GS1783 followed by kanamycin selection, and subsequent second recombination under 1% L(+)arabinose (Sigma-Aldrich)-induced I-SceI expression. Colonies were verified by PCR of the mutated region followed by sequence analysis (Macrogen), pulsed-field gel electrophoresis and restriction fragment length polymorphism. For this purpose, bacmid DNA was isolated by standard alkaline lysis from 5ml liquid cultures. Subsequently, the integrity of bacmid DNA was analyzed by digestion with restriction enzyme *Xho*I and separation in 1% PFGE agarose (Bio-Rad) gels and 0.5×TBE buffer by pulsed-field gel electrophoresis at 6 V/cm, 120-degree field angle, switch time linearly ramped from 1s to 5s over 16 h (CHEF DR III, Bio-Rad). Infectious RRV-YFP recombinants were generated as described previously (23). In short, bacmid DNA (NucleoBond Xtra Midi) was transfected into 293T cells using GenJet Ver. II (Signagen) according to manufacturer’s instructions. Transfected 293T cells were transferred onto a confluent rhesus monkey fibroblasts monolayer two days after transfection and co-cultivated until a visible cytopathic effect (CPE) was observed. For virus stocks preparations, confluent primary rhesus monkey fibroblasts were inoculated with infectious supernatant of 293T/rhesus monkey fibroblast co-cultures. After multiple rounds of replication virus-containing RF supernatant was clarified by centrifugation (4750g, 10 minutes), concentrated by overnight centrifugation (4200rpm, 4°C) and careful aspiration of approximately 95% of the supernatant. The pellet was resuspended overnight in the remaining liquid. Stocks of wt and recombinant viruses were aliquoted and stored at −80°C. Mutations were verified by PCR amplification of the respective region followed by sequence analysis (Macrogen).

### Plasmids

The pcDNA4 vector containing full-length EphB3 (ref|BC052968|, pcDNA-EphB3-myc), pcDNA6aV5 vectors containing RRV/KSHV gH and gL coding sequences (ref|GQ994935.1|, pcDNA6aV5-KSHV-gH, pcDNA3.1-KSHV-gL-Flag (14); ref|AF210726.1|, pcDNA6aV5-RRV-26-95-gH, pcDNA3.1-RRV-26-95-gL-Flag (15, 48); ref|AF083501.3|, pcDNA6aV5-RRV-17577-gH, pcDNA3.1-RRV-17577-gL-Flag (15)) were described elsewhere. RRV 26-95/ RRV 17577 recombinant gH constructs were generated based on pcDNA6aV5-RRV-26-95-gH or pcDNA6aV5-RRV-17577-gH, respectively, using ‘Round the Horn Site-directed mutagenesis. Expression plasmids pcDNA4-hPlxdc1-myc (Homo sapiens, full-length, ref |NM_020405.5|) and pcDNA4-hPlxdc2-myc (Homo sapiens, full-length, ref |NM_032812.9|) were generated by PCR based restriction cloning. The soluble ectodomain of human Plxdc2 (amino acids 31-453) without signal peptide was inserted behind a heterologous signal peptide of murine IgG-kappa into pAB61Strep by PCR based restriction cloning, resulting in a C-terminally fused IgG1 Fc-fusion protein with a C-terminal tandem Strep-Tag (pPlxdc2-FcStrep) as described previously (49). pcDNA6 vectors constructs containing human Plxdc1 ectodomain (aa 1-425) with a 6XHis Tag (pcDNA6-ectoPlxdc1-6XHis) or human Plxdc2 ectodomain (aa 1-453) with a 6XHis Tag (pcDNA6-ectoPlxdc2-6XHis) were based on pcDNA4-hPlxdc1-myc/ pcDNA4-hPlxdc2-myc, which were PCR-amplified and ligated using exonuclease-based Gibson-assembly (Gibson Assembly Mastermix, New England Biolabs). Expression plasmids pcDNA4-mmPlxdc1-myc (macaca mulatta, full-length, ref |XM_028836436.1|) and pcDNAmmPlxdc2-myc (macaca mulatta, full-length, ref |XM_028826043.1|) were generated based on PCR-amplified gBlock gene fragments (purchased from IDT) of the regions varying from human Plxdc1 (nt1-1200) and Plxdc2 (nt1-1401), respectively, and ligated in the respective PCR-amplified backbone (pcDNA4hPlxdc1-myc, pcDNA4hPlxdc2-myc) by exonuclease-based Gibson. pLenti CMV Blast DEST (706-1) (a gift from Eric Campeau & Paul Kaufman (Addgene plasmid #17451)) constructs carrying a human Plxdc1-TwinStrep or Plxdc2-TwinStrep expression cassette (pLenti-CMV-Blast-Plxdc1-Strep/ pLenti-CMV-Blast-Plxdc2-Strep) were based on pcDNA4-hPlxdc1-myc/ pcDNA4-hPlxdc2-myc, which were PCR-amplified and ligated using exonuclease-based Gibson-assembly. pLenti-CMV-Blast-EphA7-Strep was described before (20). (See S1 Table for a complete list of primers and constructs).

### Recombinant proteins

Recombinant, soluble FcStrep and Plxdc2-FcStrep-fusion proteins were purified under native conditions by Strep-Tactin chromatography from 293T cell culture supernatant. 293T cells were transfected with pAB61Strep or pPlxdc2-FcStrep using Polyethylenimine "Max" (PEI) (Polysciences) (50) as described before (20) with pAB61Strep or pPlxdc2-FcS. The protein-containing cell culture supernatant was filtered through 0.22μm PES membranes (Millipore) and passed over 0.5ml of a Strep-Tactin Superflow (IBA Lifesciences) matrix in a gravity flow Omniprep column (BioRad). Bound protein was washed with approximately 50ml phosphate buffered saline pH 7.4 (PBS) and eluted in 1ml fractions with 3mM desthiobiotin (Sigma-Aldrich) in PBS. Recombinant Plxcd1 ectodomain or Plxdc2 ectodomain protein was purified under native conditions by Ni-NTA chromatography from 293T cell culture supernatant. 293T cells were transfected pcDNA6-ectoPlxdc1-6XHis or pcDNA6-ectoPlxdc2-6XHis using PEI transfectio. The protein-containing cell culture supernatant was filtered through 0.22μm PES membranes (Millipore), concentrated using VIVAFLOW 50R (Sartorius) and passed over 1ml of a Ni-NTA Agarose (Macherey-Nagel) matrix in a gravity flow Omniprep column (BioRad). Bound protein was washed with approximately 50ml TBS (150 mM NaCl, 50 mM Tris-HCl, pH 7.6) and eluted in 1ml fractions with 500 mM Imidazole in TBS. Protein-containing fractions were pooled, concentrated via VivaSpin columns (Sartorius) and applied over Äkta Avant (GE) on a HiPrep 16/60 Sephacryl S300HR column (GE). For all recombinant proteins,protein-containing fractions were pooled and buffer exchange to PBS via VivaSpin columns (Sartorius) was performed. Protein concentration was determined by absorbance at 280nm. Aliquots were frozen and stored at −80°C. Recombinant, human, soluble EphB3-Fc (5667-B3-050) was purchased from R&D Systems.

### Lentivirus production and transduction

For production of lentiviral particles, 10cm cell culture grade petri dishes of approximately 80% confluent 293T cells were transfected with 1.4μg pMD2.G (VSV-G envelope expressing plasmid, a gift from Didier Trono (Addgene plasmid #12259), 3.6μg psPAX2 (Gag-Pol expression construct, a gift from Didier Trono (Addgene plasmid #12260), and 5μg of lentiviral expression constructs (pLenti CMV Blast DEST (706-1), pLenti-CMV-Blast-EphA7-Strep, pLenti-CMV-Blast-Plxdc1-Strep, pLenti-CMV-Blast-Plxdc2-Strep) using PEI as described before (20). The supernatant containing the pseudotyped lentiviral particles was harvested 2 to 3 days after transfection and filtered through 0.45μm CA membranes (Millipore). For transduction, lentivirus stocks were used at a 1:5 dilution unless stated otherwise. After 48h, the selection antibiotic blasticidin (Invivogen) was added to a final concentration of 10μg/ml. After initial selection the blasticidin concentration was reduced to 5μg/ml.

### Quantitative realtime-PCR-based viral genome copy number analysis and virus attachment assay

Concentrated virus samples were treated with DNAseI (0.1 units/μl) to remove any non-encapsidated DNA (37°C, overnight). Subsequently, DNAseI was deactivated and viral capsids were disrupted by heating the samples to 95°C for 30 minutes. Realtime-PCR (qPCR) was performed on a StepOne Plus cycler (Thermo Fisher Scientific) in 20μl reactions using the SensiFAST Probe Hi-ROX Kit (Bioline) (cycling conditions: 3min initial denaturation at 95°C, 40 cycles 95°C for 10s and 60°C for 35s). All primer-probe sets were purchased from IDT as complete PrimeTime qPCR Assays (primer:probe ratio = 4:1). Samples were analyzed in technical triplicates. A series of five 10-fold dilutions of bacmid DNA was used as standard for absolute quantification of viral genome copies based on qPCR of ORF73 for RRV (see S1 Table for a complete list of primers). For virus attachment assays transduced Raji cells were incubated with ice-cold virus dilutions at the indicated concentrations, normalized to genomes per cell, at 4°C for 30min. After three washes with ice-cold PBS genomic DNA was isolated using the ISOLATE II Genomic DNA Kit (Bioline) according to manufacturer’s instructions. Genome copies of used input virus preparations were determined after overnight DNAseI digest as described above. Relative values of bound viral genomes to cellular DNA were calculated on the basis of ΔCt values for viral genomic loci (ORF73 for RRV) and a cellular genomic locus (CCR5) using the same conditions described above. For attachment assays all samples were analyzed in technical duplicates.

### Infection assays, blocking experiments and flow cytometry

For infection assays cells were plated at 50 000 cells/cm^2^ (SLK, HaCaT, Raji), 25 000 cells/cm^2^ (RF) or 200 000 cells/ml (Raji, MFB5487, MMB1845) respectively. One day after plating (for adherent cells lines) or directly after plating (for suspension cell lines), cells were infected with the indicated amounts of virus. Adherent cells lines were harvested 24h post infection by brief trypsinization, followed by addition of 5% FCS in PBS to inhibit trypsin activity. Suspension cell lines were harvested 48h post infection (24h post infection for transduced Raji cells) by pipetting. Subsequently, cells were pelleted by centrifugation (1200rpm, 10min), washed once with PBS, re-pelleted and fixed in PBS supplemented with 4% formaldehyde (Carl Roth). Block of RRV infection with soluble decoy receptor was assayed by infection with virus inocula that were pre-incubated with the indicated concentrations of soluble EphB3-Fc, hPlxdc1-Fc, hPlxdc2-Fc or Fc alone at room temperature for 30min. Calculation of molarity was based on diametric proteins. Cell harvest and preparation for flow cytometry analyses was performed as described above. A minimum of 5 000 – 10 000 cells was analyzed per sample for YFP expression on a LSRII flow cytometer (BD Biosciences). Data was analyzed using Flowing Software (Version 2.5).

### Immunoprecipitation and Western blot analysis

For interaction analysis of gH-V5/gL-Flag complexes with Plxdc1/2, 293T cells were transfected using PEI as described before. Lysates of 293T cells transfected with the respective expression constructs for recombination gH-V5/gL-Flag complexes were prepared in NP40 lysis buffer (1% Nonidet P40 Substitute (Sigma-Aldrich), 150mM NaCl (Sigma-Aldrich), 50mM HEPES (VWR), 1mM EDTA (Amresco) with freshly added Protease Inhibitor Cocktail, General Use (Amresco)) and protein content was determined by Bradford assay using Roti-Quant (Roth) according to manufacturer’s instructions. 20μg total protein was denatured in 1× SDS sample buffer (Morris formulation) at 95°C for 5min, separated by polyacrylamide gel electrophoresis (PAGE) using 8–16% Tris-Glycine polyacrylamide gradient gels (Thermo Fisher Scientific) with Tris-Glycine SDS running buffer (25mM Tris, 192mM glycine, 0.1% SDS) and transferred to 0.45μm (0.22μm for blots containing gL) Polyvinylidendifluorid (PVDF) membranes (200mA/Gel, max 30V, 1h in Towbin buffer (25mM Tris, 192mM glycine) with 20% methanol) in a wet tank system (Mini Blot Module, Thermo Fisher). The membranes were blocked in 5% dry milk powder in TBS-T (5mM Tris, 15mM NaCl, 0.05% Tween20) for 1h, at room temperature, washed once in TBS-T and incubated with the respective antibodies for 2h at room temperature or overnight at 4°C (see S1 Table for a complete list of antibodies). After three washes with TBS-T, the membranes were incubated with the respective HRP-conjugated secondary antibody in 5% dry milk powder in TBS-T for 1h at room temperature washed three times in TBS-T and imaged on an ECL ChemoCam 3.2 Imager (Intas) using Immobilon Forte Western HRP substrate (Merck Millipore). For pulldown of gH-V5 or gH-V5/ gL-Flag complexes, the amount of input lysate between wt and mutant gH constructs was normalized to gH expression as determined by Western blot and diluted to equal volume with cell lysate from non-transfected 293T cells prior to immunoprecipitation. Subsequently, lysates were incubated with 0.5μg V5-tag antibody (Bio-Rad) and ProteinG sepharose (GenScript) overnight at 4°C with agitation. After three washes in NP40 lysis buffer, ProteinG beads with pre-coupled complexes were incubated overnight at 4°C with agitation with lysate of full-length human or macaca mulatta Plxdc1-myc or Plxdc2-myc expression plasmid transfected 293T cells normalized to Plxdc expression. Volumes were adjusted with lysate from untransfected 293T cells. ProteinG beads were collected by brief centrifugation and washed 3 times in NP40 lysis buffer. Precipitates were heated in 2x SDS sample buffer (95°C, 5min) and analyzed by Western blot as described above. For co-immunoprecipitation of soluble Plxdc1/2-Strep constructs with gH-V5/gL-Flag and EphB3-myc, supernatant of Plxdc1/2-FcStrep transfected 293T cells was incubated with StrepTactinXT beads (IBA) overnight at 4°C with agitation. After three washes in NP40 lysis buffer, StrepTactinXT beads with pre-coupled Plxdc1/2-Strep were incubated overnight at 4°C with agitation with equal amounts of lysate of full-length human EphB3-myc, RRV 26-95 gH-V5/gL-Flag or 17577 gH-V5/gL-Flag expression plasmid transfected 293T cells or the indicated combinations. Volumes were adjusted with lysate of untransfected 293T cells.

### Enzyme-linked immunosorbent assay (ELISA)

F96 Maxisorp Nunc-Immuno Plates (Thermo Fisher Scientific) were coated with recombinant RRV 26-95 gH-FcStrep/gL (described previously (14)) at 1μg/ml in PBS overnight. After three washes with PBS-T, the wells were blocked with 10% FBS in PBS for 2h. Incubation with Plxdc1 ectodomain or Plxdc2 ectodomain was performed for 2h at room temperature in 10% FBS in PBS. The plates were washed three times with PBS-T. Bound protein was detected via the C-terminal 6XHis Tag using 6XHis Antibody MA1-135 (Invitrogen) followed by three washes in TBS-T and incubation with donkey anti-mouse horseradish peroxidase (HRP)-coupled secondary antibody (Dianova). After three washes, 3,3ʹ,5,5ʹ-Tetramethylbenzidin (TMB) substrate (Thermo Fisher Scientific) was added and the reaction was stopped by adding 1M HCl. The plates were imaged on a Biotek Synergy 2 plate reader.

### Structure prediction and analysis

Homology based structure prediction was performed using the Iterative Threading ASSembly Refinement (I-TASSER) server on standard settings for structure prediction of RRV 26-95 gH and gL based on the crystal structure of the EBV gH/gL complex (3PHF). Modeling of the RRV 26-95 gH/gL complex was additionally performed using both the SPRING and CO-THreader algorithms for protein-protein complex structure and multi-chain protein threading with no differences between determined structures. Resulting I-TASSER structures were aligned to the gH/gL CO-THreader model with the VMD 1.9.3 OpenGL RMSD Trajectory Tool based on amino acids 25 to 62 of gH (RMSD of 0.344Å) and amino acids 2 to 100 of gL (RMSD of 2.697Å) to generate the depicted model. All further analyses and visualizations were performed using VMD 1.9.3 OpenGL.

### Mathematical and statistical analysis

Statistical difference between groups was determined by unpaired Student’s t-tests followed by Bonferroni correction for multiple comparisons. All Statistical analyses were performed with GraphPad Prism version 6. For all statistics, *: p-value < 0.05, **: p-value < 0.01, ***: p-value < 0.001, ns: not significant.

## Supporting information

S1 Table List of primers and antibodies

## ACKNOWLEDGMENTS

This work was supported by grants to A.S.H. from the Deutsche Forschungsgemeinschaft (HA 6013/1, HA 6013/4-1), the Wilhelm-Sander Foundation (project 2019.027.1), and the Interdisciplinary Center for Clinical Research Erlangen (IZKF, grant J44), by the Deutsche Forschungsgemeinschaft, grants CRC 796, TP B1 to A.E., by grant RO1 AI072004 from the National Institutes of Health (NIH) to R.C.D., and by base grant RR00168 from the NIH to the New England Primate Research Center.

---

**S1 Table. List of primers, and antibodies used in this study**.

